## Supporting information for "Thalamocortical contributions to hierarchical cognitive control"

### S1. Behavioral results considering response repetition and cue repetition

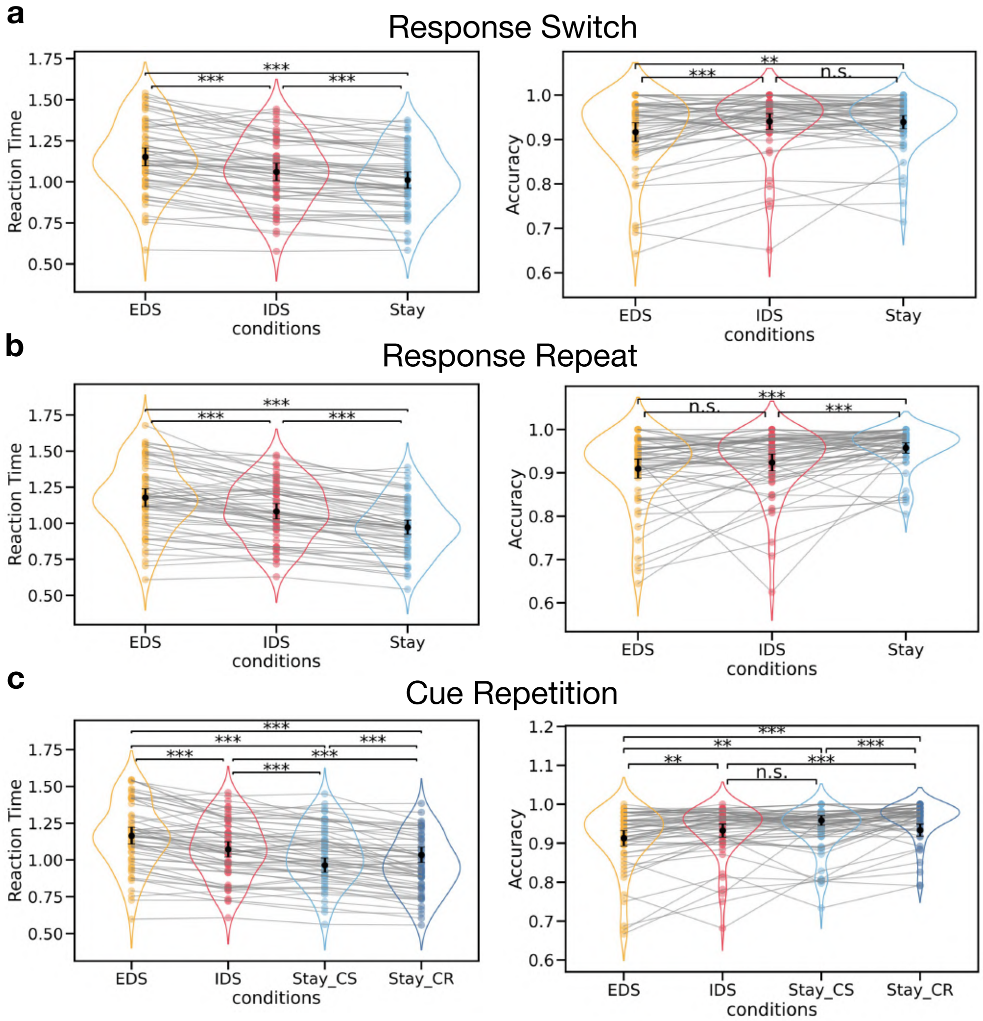


***S1. Behavioral results considering response repetition and cue repetition.*** *(a) Behavioral results for response switching. (b) Behavioral results for response repetition. (c) Behavioral results for cue repetition. (a)(b)(c) the left panel is reaction time, and the right panel is accuracy. *** p<0.001. ** p<0.01. n.s. non-significant. The error bar represents the 95% confidential interval. The black dot indicates the mean value, while the colored dot represents data from individual subjects. Lines connect the data points for each subject across different conditions.*

### S2. Brain regions activated during hierarchical task-switching

**Table 1: Brain regions activated during EDS**

| Voxels | CM coordinates | Regions | t-value (mean ± SEM) |
|---|---|---|---|
| 35996 | [-0.5, 65.8, -0.7] | intraparietal sulcus; lateral occipital gyrus; the fusiform gyrus; the inferior temporal gyrus; thalamus; basal ganglia | 3.72 ± 0.01 |
| 2838 | [44.6, -14.8, 32.1] | left middle frontal gyrus | 3.25 ± 0.02 |
| 1498 | [1.1, -12.2, 50.5] | dorsal medial prefrontal cortex | 3.33 ± 0.03 |
| 963 | [-47.4, -35.3, 25.2] | right superior frontal gyrus | 3.21 ± 0.03 |
| 487 | [36.7, -18.7, 3.9] | left insula | 3.58 ± 0.06 |
| 434 | [-45.5, -8.7, 28.7] | right middle frontal gyrus | 3.17 ± 0.04 |
| 433 | [-36.2, -21.5, 2.5] | right insula | 3.42 ± 0.04 |
| 273 | [-0.2, 30.6, 27.6] | posterior cingulate cortex | 3.23 ± 0.05 |
| 229 | [-37.4, -0.9, 56.3] | right precentral sulcus | 2.92 ± 0.05 |
| 186 | [0.9, -5.9, 83.0] | left superior frontal gyrus | 2.90 ± 0.05 |
| 154 | [11.7, 74.7, 10.4] | left cuneus | 2.77 ± 0.05 |
| 132 | [44.1, 3.0, 10.8] | left precentral sulcus | 2.97 ± 0.07 |
| 101 | [23.6, 51.4, 24.6] | left precuneus | 2.75 ± 0.06 |
| 101 | [60.4, 20.6, 26.9] | left postcentral sulcus | 2.65 ± 0.05 |

**Table 2: Brain regions activated during IDS**

| Voxels | CM coordinates | Regions | t-value (mean ± SEM) |
|---|---|---|---|
| 28981 | [-1.4, 67.5, -0.7] | intraparietal sulcus; lateral occipital gyrus; the fusiform gyrus; the inferior temporal gyrus; thalamus; basal ganglia | 3.67 ± 0.01 |
| 1277 | [47.8, -13.6, 30.4] | left middle frontal gyrus | 3.13 ± 0.03 |
| 964 | [2.2, -9.8, 51.4] | dorsal medial prefrontal cortex | 3.14 ± 0.03 |
| 502 | [-47, -31.7, 23.7] | right superior frontal gyrus | 3.20 ± 0.04 |
| 323 | [30.4, 5.3, 53.5] | left precentral sulcus | 3.17 ± 0.05 |
| 286 | [35.6, -18.6, 6.6] | left insula | 3.22 ± 0.05 |
| 278 | [-45.0, -8.0, 28.4] | right middle frontal gyrus | 3.26 ± 0.06 |
| 249 | [-35.8, -21.5, 4.4] | right insula | 2.93 ± 0.04 |
| 174 | [59.4, 20.0, 25.7] | left postcentral sulcus | 2.67 ± 0.04 |
| 163 | [3.3, -14.8, 79.6] | left superior frontal gyrus | 2.84 ± 0.05 |
| 155 | [44.1, 2.9, 13.1] | left precentral sulcus | 3.16 ± 0.08 |
| 101 | [-37.0, 0.4, 56.0] | right precntral sulcus | 2.84 ± 0.07 |
| 68 | [-45.5, -50.3, 23.1] | right superior frontal gyrus | 2.89 ± 0.10 |
| 60 | [39.4, 23.3, 57.2] | left precentral sulcus | 2.94 ± 0.10 |

**Table 3: Brain regions activated during Stay**

| Voxels | CM coordinates | Regions | t-value (mean ± SEM) |
|---|---|---|---|
| 26416 | [-1.6, 67.0, -1.9] | intraparietal sulcus; lateral occipital gyrus; the fusiform gyrus; the inferior temporal gyrus; thalamus; basal ganglia | 3.71 ± 0.01 |
| 862 | [2.6, -9.7, 51.7] | dorsal medial prefrontal cortex | 3.07 ± 0.03 |
| 807 | [46.3, -10.6, 30.3] | left middle frontal gyrus | 3.07 ± 0.03 |
| 380 | [-46.5, -31.1, 23.3] | right superior frontal gyrus | 3.22 ± 0.05 |
| 251 | [35.1, -18.3, 7.9] | left insula | 3.15 ± 0.06 |
| 251 | [-44.3, -7.9, 28.3] | right superior frontal gyrus | 2.87 ± 0.04 |
| 224 | [-34.7, -21.6, 6.1] | right insula | 2.89 ± 0.05 |
| 195 | [44.1, 2.4, 13.2] | left insula | 3.36 ± 0.08 |
| 181 | [31.5, 5.6, 52.3] | left precentral sulcus | 2.93 ± 0.05 |
| 141 | [59.0, 20.0, 25.3] | left postcentral sulcus | 2.58 ± 0.04 |
| 104 | [1.4, -12.2, 80.8] | left superior frontal gyrus | 2.84 ± 0.06 |
| 78 | [-36.9, 0.6, 55.0] | right precentral sulcus | 2.79 ± 0.07 |
| 60 | [39.4, 20.7, 57.5] | left precentral sulcus | 2.90 ± 0.10 |

**Table 4: Brain regions activated during EDS-Stay**

| Voxels | CM coordinates | Regions | t-value (mean ± SEM) |
|---|---|---|---|
| 46163 | [5.0, 29.5, 17.5] | superior and middle frontal gyrus; insula; posterior cingulate gyrus; cuneus; intraparietal sulcus; left thalamus; left basal ganglia | 3.05 ± 0.004 |
| 536 | [-12.3, 1.3, 14.7] | right thalamus; right basal ganglia | 2.89 ± 0.03 |
| 244 | [53.7, 20.5, 11.2] | left superior temporal gyrus | 2.73 ± 0.04 |
| 203 | [-55.8, 13.9, 7.4] | right superior temporal gyrus | 2.85 ± 0.05 |
| 163 | [-50.7, 30.9, -2.6] | right superior temporal sulcus | 2.64 ± 0.04 |
| 101 | [21.4, 27.5, 74.8] | left precentral sulcus | 2.70 ± 0.06 |

**Table 5: Brain regions activated during EDS-IDS**

| Voxels | CM coordinates | Regions | t-value (mean ± SEM) |
|---|---|---|---|
| 5566 | [6.8, 62.8, 47.8] | intraparietal sulcus | 2.88 ± 0.01 |
| 3948 | [42.7, -25.6, 22.6] | left middle frontal gyrus; left superior frontal gyrus; left insula | 2.87 ± 0.01 |
| 1832 | [-43.6, -33.6, 22.4] | right middle frontal gyrus; right superior frontal gyrus; right insula | 2.78 ± 0.01 |
| 1790 | [0.3, -20.7, 51.8] | dorsal medial prefrontal cortex | 2.80 ± 0.01 |
| 613 | [0, 25.2, 31.6] | posterior cingulate cortex | 2.85 ± 0.03 |
| 496 | [7.2, 7.6, 11.0] | thalamus; left basal ganglia | 2.66 ± 0.02 |
| 493 | [-2.6, 74.8, 8.2] | cuneus | 2.67 ± 0.02 |
| 448 | [51.9, 68.0, -21.5] | inferior temporal gyrus | 2.71 ± 0.03 |
| 437 | [-40.2, -20.2, 0.3] | right insula | 2.74 ± 0.03 |
| 273 | [62.2, 36.4, -5.6] | left middle temporal gyrus | 2.59 ± 0.03 |

| | | | |
|---|---|---|---|
| 263 | [-14.5, -5.4, 11.2] | right basal ganglia | 2.67± 0.03 |
| 5566 | [6.8, 62.8, 47.8] | intraparietal sulcus | 2.88 ± 0.01 |
| 3948 | [42.7, -25.6, 22.6] | left middle frontal gyrus; left superior frontal gyrus; left insula | 2.87 ± 0.01 |

**Table 6: Brain regions activated during IDS-Stay**

| Voxels | CM coordinates | Regions | t-value (mean ± SEM) |
|---|---|---|---|
| 2526 | [20.6, 62.6, 56.7] | left intraparietal sulcus | 2.71 ± 0.01 |
| 735 | [52.2, -17.9, 30.5] | left caudal middle frontal sulcus | 2.68 ± 0.02 |
| 523 | [28.8, 3.4, 62.6] | left precentral sulcus | 2.80 ± 0.03 |
| 270 | [-55.5, 13.8, 7.7] | right superior temporal sulcus | 2.71 ± 0.03 |
| 252 | [10.9, -11.3, 74.7] | left caudal superior frontal sulcus | 2.72 ± 0.04 |
| 200 | [2.6, -9.4, 52.8] | left dorsal medial prefrontal cortex | 2.64 ± 0.03 |
| 177 | [56.5, 21.0, 11.3] | left superior temporal sulcus | 2.61 ± 0.04 |
| 164 | [30.3, 40.3, 73.4] | left postcentral sulcus | 2.75 ± 0.05 |
| 153 | [-29.9, 2.0, 60.7] | right caudal middle frontal sulcus | 2.77 ± 0.05 |
| 114 | [6.0, 21.9, 86.2] | left precentral sulcus | 2.87 ± 0.06 |
| 96 | [54.0, 69.2, -13.8] | left inferior temporal gyrus | 2.89 ± 0.07 |
| 93 | [-48..7, -16.2, -4,1] | right insula | 2.69 ± 0.08 |
| 91 | [-60.2, 24.8, -38.9] | right inferior temporal gyrus | 2.97 ± 0.07 |
| 74 | [-61.3, 61.8, -11.5] | right inferior temporal gyrus | 2.69 ± 0.08 |
| 60 | [-37.0, 44.2, 39.9] | right intraparietal sulcus | 2.60 ± 0.07 |

**Table 7: Brain regions activated during EDS_RR-Stay_RR**

| Voxels | CM coordinates | Regions | t-value (mean ± SEM) |
|---|---|---|---|
| 5821 | [34.6, 63.7, 54.0] | intraparietal sulcus | 2.86 ± 0.01 |
| 3331 | [44.9, -37.7, 14.0] | left middle frontal gyrus | 2.91 ± 0.01 |
| 1148 | [2.0, 1.9, 64.0] | dorsal medial prefrontal cortex | 2.81 ± 0.02 |
| 710 | [-49.6, -46.3, 1.5] | right middle frontal gyrus | 2.73 ± 0.02 |
| 578 | [48.4, -15.3, -6] | left insula | 2.91 ± 0.03 |
| 419 | [-34.1, -18.8, 6.5] | right insula | 2.75 ± 0.03 |
| 256 | [-32.4, -1.6, 66.5] | right caudal middle frontal gyrus | 2.78 ± 0.04 |
| 233 | [2.0, 34.5, 26.5] | posterior cingulate cortex | 2.88 ± 0.05 |
| 197 | [-13.5, 0.1, 21.5] | right thalamus, right basal ganglia | 2.64 ± 0.04 |
| 167 | [19.1, 22.5, 24.0] | left thalamus, left basal ganglia | 2.64 ± 0.04 |
| 145 | [-10.1, -29.1, 29.0] | right anterior cingulate cortex | 2.59 ± 0.04 |
| 131 | [57.0, 24.2, -43.5] | left inferior temporal gyrus | 2.98 ± 0.06 |
| 94 | [14.0, -5.0, 11.5] | left basal ganglia | 2.57 ± 0.06 |
| 93 | [3.7, 20.8, 11.5] | medial thalamus | 2.67 ± 0.05 |

**Table 8: Brain regions activated during EDS_RR-IDS_RR**

| Voxels | CM coordinates | Regions | t-value (mean ± SEM) |
|---|---|---|---|
| 1342 | [38.0, 58.6, 49.0] | left intraparietal sulcus | 2.78 ± 0.02 |
| 1081 | [58.7, -18.8, 31.5] | left caudal middle frontal sulcus | 2.72 ± 0.02 |
| 1067 | [-8.4, 75.8, 69.0] | precuneus | 2.76 ± 0.02 |
| 779 | [39.8, -44.6, 4.0] | left rostral middle frontal sulcus | 2.71 ± 0.02 |
| 777 | [-37.6, 67.2, 66.5] | right intraparietal sulcus | 2.72 ± 0.02 |
| 604 | [3.7, -15.3, 51.5] | dorsal medial prefrontal cortex | 2.66 ± 0.02 |
| 259 | [-44.5, -34.2, 39.0] | right middle frontal gyrus | 2.62 ± 0.03 |
| 254 | [34.6, -20.5, 1.5] | left insula | 2.81 ± 0.04 |
| 205 | [29.5, -13.6, 61.5] | left superior frontal gyrus | 2.68 ± 0.04 |
| 198 | [2.0, 36.2, 34.0] | posterior cingulate cortex | 2.67 ± 0.04 |
| 197 | [-51.3, -10.2, 26.5] | right precentral sulcus | 2.63 ± 0.04 |
| 167 | [-49.6, -29.1, 21.5] | right rostral middle frontal sulcus | 2.71 ± 0.04 |
| 166 | [2.0, 34.5, 6.5] | medial thalamus | 2.60 ± 0.04 |
| 162 | [-34.1, -22.2, 9.0] | right insula | 2.58 ± 0.03 |
| 142 | [5.4, -3.3, 79.0] | left superior frontal gyrus | 2.79 ± 0.05 |

**Table 9: Brain regions activated during IDS_RR-Stay_RR**

| Voxels | CM coordinates | Regions | t-value (mean ± SEM) |
|---|---|---|---|
| 1494 | [36.3, 70.6, 54.0] | left intraparietal sulcus | 2.73 ± 0.02 |
| 600 | [0.2, -29.1, 36.5] | dorsal medial prefrontal cortex | 2.58 ± 0.02 |
| 435 | [36.3, -49.7, 4.0] | left rostral middle frontal sulcus | 2.64 ± 0.03 |
| 411 | [57.0, -18.8, 31.5] | left caudal middle frontal sulcus | 2.68 ± 0.03 |
| 365 | [-35.9, 67.2, 46.5] | right intraparietal sulcus | 2.61 ± 0.03 |
| 299 | [-13.5, 101.6, -13.5] | right occipital cortex | 2.82 ± 0.03 |
| 217 | [44.9, -17.1, 6.5] | left insula | 2.73 ± 0.04 |
| 147 | [-35.9, -22.2, 1.5] | right insula | 2.57 ± 0.03 |
| 119 | [-11.8, 70.6, 39.0] | right parieto-occipital sulcus | 2.60 ± 0.05 |
| 116 | [31.2, 3.6, 64.0] | left precentral sulcus | 2.73 ± 0.05 |
| 106 | [-44.5, 43.1, 46.5] | right intraparietal sulcus | 2.58 ± 0.04 |
| 103 | [48.4, 39.7, -1.0] | left middle temporal sulcus | 2.53 ± 0.04 |
| 102 | [3.7, 41.4, 24.0] | posterior cingulate cortex | 2.67 ± 0.06 |
| 96 | [-49.6, -30.8, 34.0] | right rostral middle frontal gyrus | 2.61 ± 0.05 |
| 80 | [39.8, -30.8, 19] | left middle frontal gyrus | 2.64 ± 0.05 |

**Table 10: Brain regions activated during EDS-Stay_CR**

| Voxels | CM coordinates | Regions | t-value (mean ± SEM) |
|---|---|---|---|
| 7339 | [32.9, 68.9, 46.5] | intraparietal sulcus | 3.07 ± 0.01 |
| 6763 | [41.5, 0.1, 36.5] | left middle frontal sulcus | 3.03 ± 0.01 |
| 1997 | [-53.1, -11.9, 26.5] | right middle frontal sulcus | 2.91 ± 0.02 |
| 393 | [7.1, 72.3, 11.5] | occipital cortex | 2.70 ± 0.03 |
| 379 | [2.0, 31.1, 29.0] | posterior cingulate cortex | 3.06 ± 0.04 |

| | | | |
|---|---|---|---|
| 378 | [8.8, 3.6, 14.0] | left thalamus; left basal ganglia | 2.83 ± 0.04 |
| 359 | [-46.2, -17.1, -6.0] | right insula | 2.87 ± 0.04 |
| 266 | [-11.8, 1.9, 16.5] | right thalamus; right basal ganglia | 2.78 ± 0.04 |
| 148 | [-44.5, -53.2, 6.5] | right rostral middle frontal sulcus | 2.61 ± 0.04 |

**Table 11: Brain regions activated during IDS-Stay_CR**

| Voxels | CM coordinates | Regions | t-value (mean ± SEM) |
|---|---|---|---|
| 1413 | [32.9, 68.9, 46.5] | left intraparietal sulcus | 2.83 ± 0.02 |
| 648 | [51.8, -8.5, 31.5] | left middle frontal sulcus | 2.73 ± 0.02 |
| 363 | [32.9, 5.3, 61.5] | left precentral sulcus | 2.80 ± 0.03 |
| 335 | [-35.9, 70.6, 64.0] | right intraparietal sulcus | 2.65 ± 0.03 |
| 260 | [0.2, -11.9, 49.0] | dorsal medial frontal cortex | 2.61 ± 0.03 |
| 174 | [29.5, -65.2, 26.5] | left rostral middle sulcus | 2.78 ± 0.05 |
| 154 | [-54.8, -11.9, 26.5] | right precentral sulcus | 2.72 ± 0.05 |
| 123 | [-46.2, 44.8, 61.5] | right intraparietal sulcus | 2.65 ± 0.05 |
| 105 | [53.5, -36.0, 24.0] | left superior frontal sulcus | 2.69 ± 0.06 |
| 87 | [-37.6, -1.6, 56.5] | right caudal middle frontal sulcus | 2.81 ± 0.06 |
| 76 | [-46.2, -32.5, 31.5] | right middle frontal sulcus | 2.88 ± 0.08 |
| 71 | [26.0, 41.4, 76.5] | left postcentral sulcus | 2.71 ± 0.06 |

**S3. Cortical evoked response to hierarchical task-switching considering response repetition and cue repetition.**

We found that, when considering the response and cue repetition, the overall cortical response patterns were consistent to our main findings. Specifically, stronger evoked responses were observed in the frontoparietal and temporal regions in response to changing task contexts (EDS_RR-Stay_RR and EDS_RR-IDS_RR for response repetition; EDS-Stay_CR, EDS-Stay_CS for cue repetition). These areas included the rostral middle frontal gyrus, the inferior frontal sulcus, the insula, the precentral sulcus, the postcentral sulcus, the intraparietal sulcus, the medial frontal gyrus, the posterior cingulate gyrus, the precuneus, the cuneus and middle temporal lobe (CM coordinates available in S2). We did not observe any significant positive clusters when comparing cue repeat for cue switch (Stay_CS-Stay_CR).

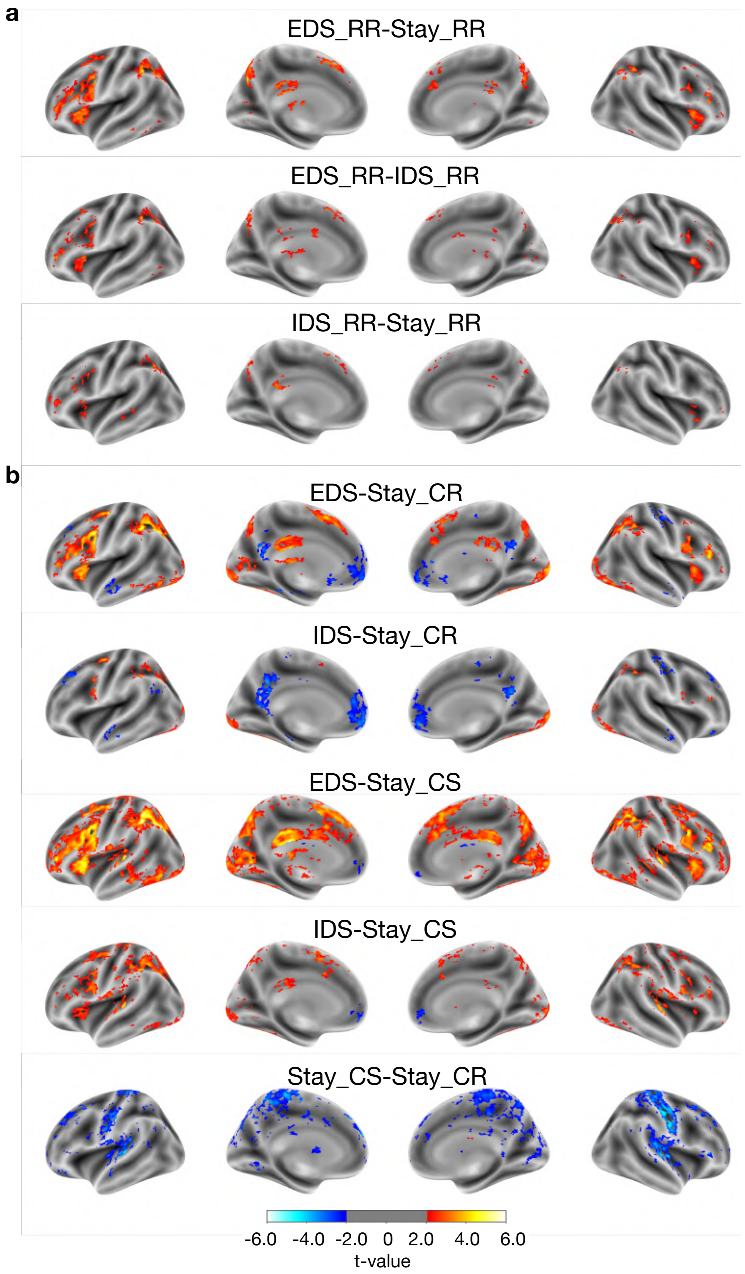


***S3. Cortical evoked response to hierarchical task-switching considering response repetition and cue repetition.*** *(a) Contrasts between hierarchical task-switching conditions, accounting for trials where the current trial's decision (choosing 'yes' or 'no') repeats from the previous trial (RR). (b)Contrasts between task-switching conditions, separate trials where the cue of the current trial either repeats (CR) or switches (CS) from the previous trial. The results were first thresholded at a voxel-level threshold of* $p < 0.05$*, followed by cluster correction procedure with a cluster level threshold of* $p<0.05$*, only showing clusters with a minimum cluster size (k) of 58 voxels. Clusters were defined as groups of voxels that are connected by sharing a face with their neighboring voxels.*

1 **S4. Hemodynamic response functions (HRF) of major thalamic nuclei**

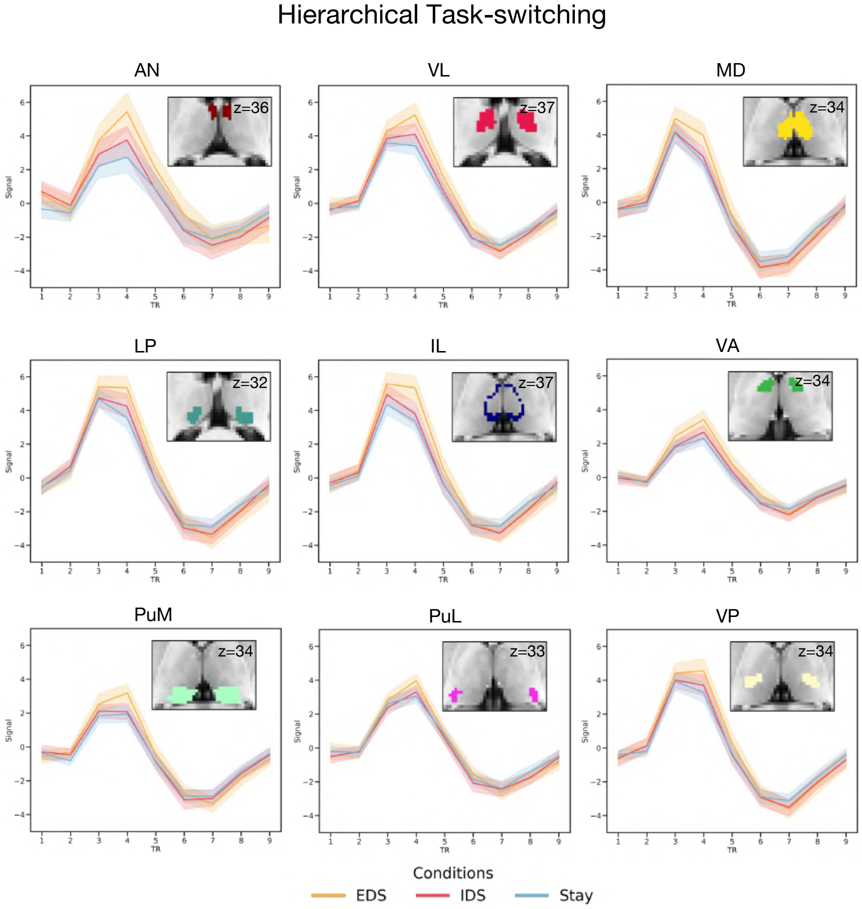


2

3 ***S4. Hemodynamic response functions of 9 major thalamic nuclei.***

**S5. Thalamic voxel-wise evoked responses to hierarchical task-switching considering response repetition and cue repetition.**

We found that, when considering the response and cue repetition, the overall thalamic response patterns were consistent to our main findings. Specifically, the anterior, ventroanterior and mediodorsal thalamic regions showed stronger evoked response when updating to higher-level task representations (EDS_RR-IDS_RR and EDS_RR-Stay_RR for response repetition; EDS-Stay_CR, EDS-Stay_CS, and IDS-Stay_CS for cue repetition). The anterior thalamic region exhibited a stronger evoked response to cue switching (Stay_CS-Stay_CR). However, no significant clusters were observed in the thalamus for IDS_RR-Stay_RR and IDS-Stay_CR conditions.

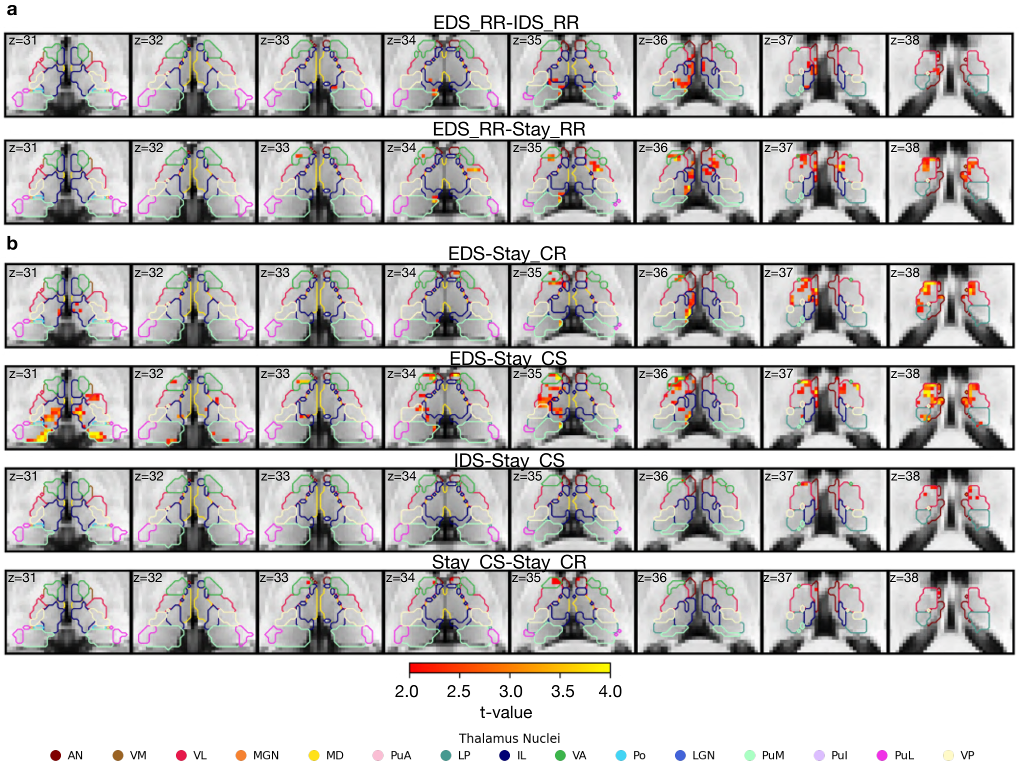


***S5. Thalamic voxel-wise evoked responses to hierarchical task-switching considering response repetition and cue repetition.*** *(a) Contrasts between conditions, trials where current trial's decision (choosing 'yes' or 'no') repeats the previous trial (RR). (b) Contrasts between task-switching conditions, separate trials where the cue of the current trial either repeats (CR) or switches (CS) from the previous trial. The results were first thresholded at a voxel-level threshold of p < 0.05, followed by cluster correction procedure with a cluster level threshold of p<0.05, only showing clusters with a minimum cluster size (k) of 58 voxels. Clusters were defined as groups of voxels that are connected by sharing a face with their neighboring voxels.*

### S6. Voxel-to-voxel thalamocortical interaction model performance

Thalamocortical interaction model outperformed two null models, which assumed that neither thalamic activity patterns nor thalamocortical functional connectivity carry task-specific information across three hierarchical task-switching conditions when predicting different levels of cortical representation ($p$<0.0001; S6a).

There was no difference in prediction accuracy between the three hierarchical task switching conditions (Context: $F(2,116)=0.41$, $p=0.66$; Context x CO: $F(2,116)=1.03$, $p=0.36$; Context x SH: $F(2,116)=1.70$, $p=0.16$; Decision: $F(2,116)=0.44$, $p=0.65$; S6b).

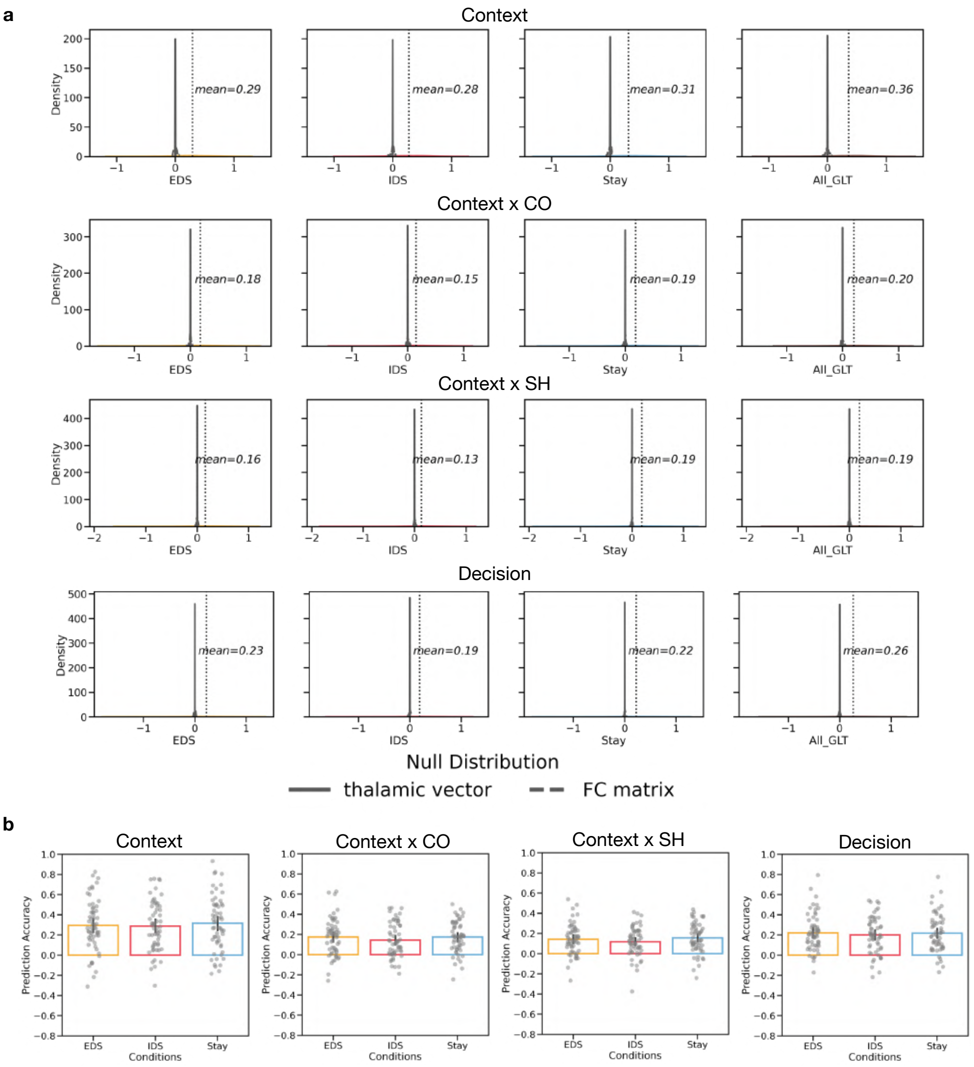


***S6. Voxel-to-voxel thalamocortical interactional model.*** *(a) Thalamocortical interaction model compared to null models; (b) Model performance of three hierarchical task switching conditions when predicting different cortical representations.*

**S7. BG voxel-wise evoked responses to hierarchical task-switching, accounting for both response and cue repetition.**

We found that, when considering the response repetition and cue repetition, the overall BG response patterns were consistent to our main findings. Specifically, the caudate activity was stronger when updating to the higher level of task representation in both response repetition (EDS_RR-Stay_RR, EDS_RR-IDS_RR, and IDS_RR-Stay_RR) and cue repetition (EDS-Stay_CR) conditions. Additionally, the putamen and globus pallidus showed increased activity during updates to higher level task representations, particularly in contrasts involving cue switching (EDS-Stay_CS and IDS-Stay_CS). No significant clusters were observed in the IDS-Stay_CR condition. Both the caudate and putamen also exhibited stronger responses to cue switching (Stay_CS-Stay_CR).

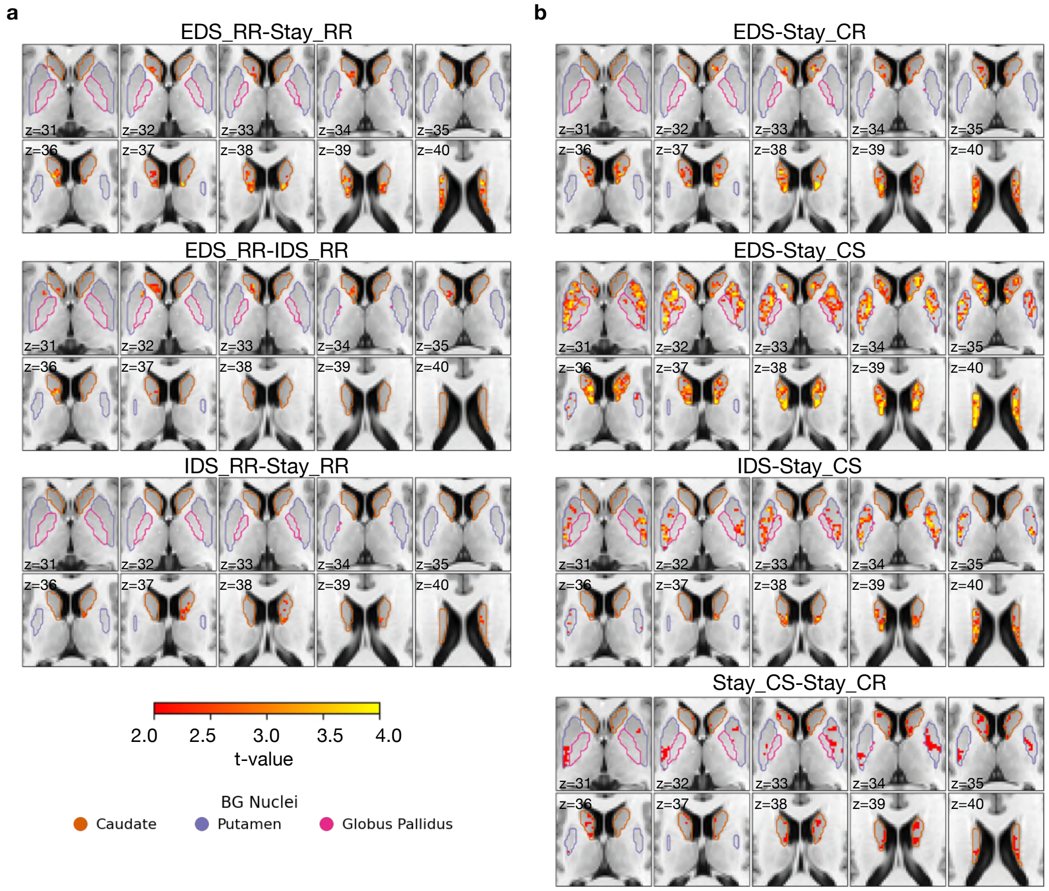


***S7. BG voxel-wise evoked response to hierarchical task-switching, accounting for both response repetition and cue repetition.*** *(a) Contrasts between conditions, separating trials where current trial's decision (choosing 'yes' or 'no') repeats from the previous trial (RR). (b) Contrasts between task-switching conditions, where the cue in the current trial either repeats (CR) or switches (CS) from the previous trial. The results were first thresholded at a voxel-level threshold of* $p < 0.05$*, followed by cluster correction procedure with a cluster level threshold of* $p<0.05$*, only showing clusters with a minimum cluster size (k) of 58 voxels. Clusters were defined as groups of voxels that are connected by sharing a face with their neighboring voxels.*

### S8. Voxel-to-voxel BG nuclei-cortical interaction model compared to null models

We found that the globus pallidus-cortical interaction model did not significantly differ from the null model in predicting the Context x Color task representation in the IDS condition ($p = 0.0838$). However, other basal ganglia (BG)-cortical interaction models outperformed the null models that assumed neither BG nuclei activity patterns nor BG-cortical functional connectivity carry task-specific information across three hierarchical task-switching conditions ($p < 0.0001$; S8).

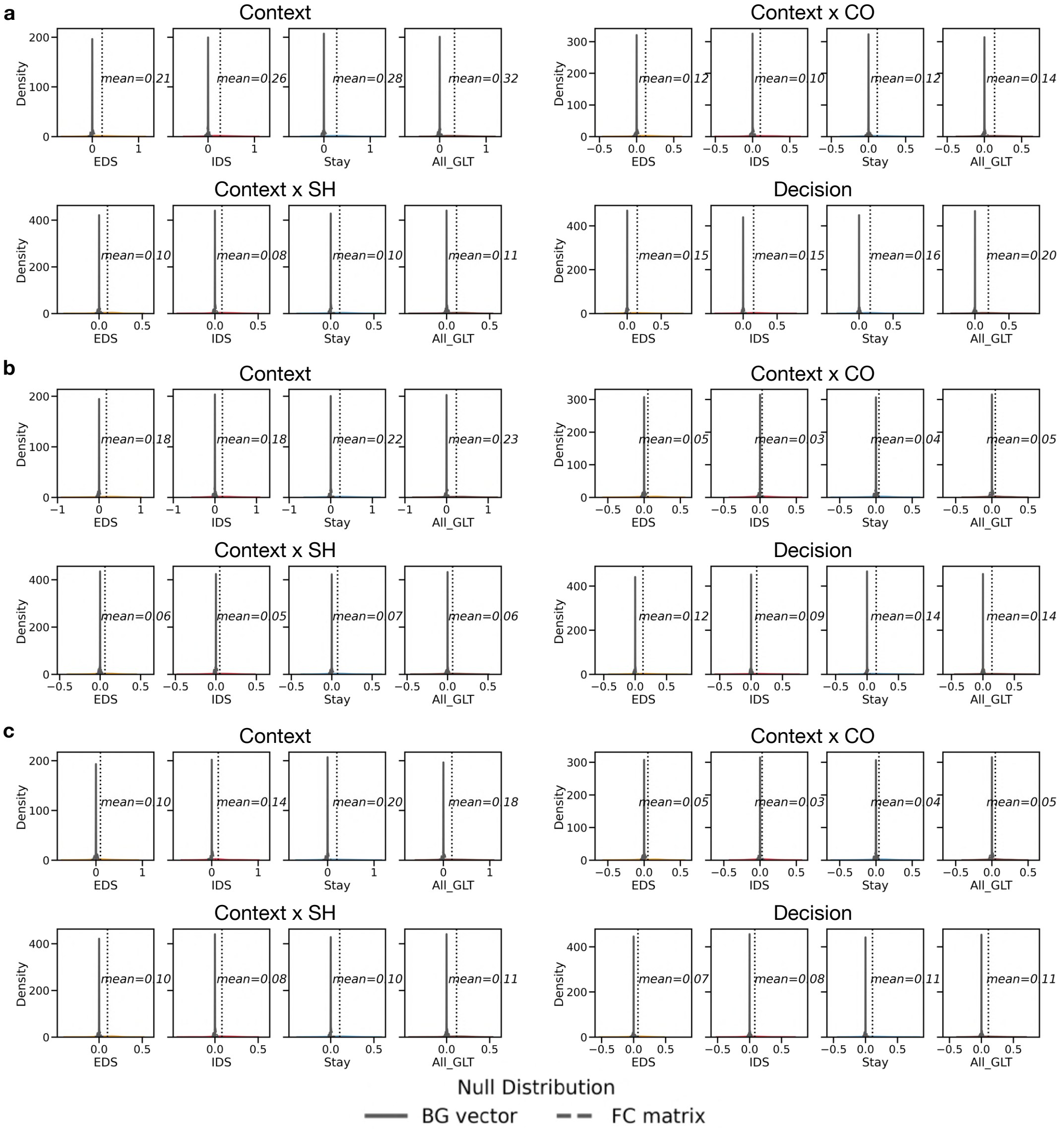


***S8. Voxel-to-voxel BG-cortical interaction model compared to two null models.*** *(a) Caudate-cortical interaction model compared to two null models; (b) Putamen-cortical interaction model compared to two null models; (c) Globus pallidus-cortical interaction model compared to two null models.*

**S9. Voxel-to-voxel BG nuclei-cortical interaction model performances for predicting hierarchical task-switching conditions and cortical representations**

We only observed significant condition difference of globus pallidus-cortical interaction model in predicting context cortical representation ($F(2,116)=5.15$, $p=0.01$, EDS vs. Stay: $t(58)=-2.82$, $p=0.02$; EDS vs. IDS: $t(58)=-1.43$, $p=0.47$; IDS vs. Stay: $t(58)=-2.03$, $p=0.14$). We did not observe any other significance difference in other models or conditions (Figure S9a; Caudate: Context: $F(2,116)=2.44$, $p=0.09$; CxCO: $F(2,116)=0.33$, $p=0.72$; CxSH: $F(2,116)=1.32$, $p=0.27$; Decision: $F(2,116)=0.35$, $p=0.71$; Putamen: Context: $F(2,116)=1.26$, $p=0.29$; CxCO: $F(2,116)=0.05$, $p=0.95$; CxSH: $F(2,116)=1.02$, $p=0.36$; Decision: $F(2,116)=1.70$, $p=0.19$; Globus pallidus: CxCO: $F(2,116)=0.49$, $p=0.61$; CxSH: $F(2,116)=1.01$, $p=0.37$; Decision: $F(2,116)=2.60$, $p=0.08$).

We found that BG-cortical interaction models display similar prediction pattern when predicting different cortical representations (Caudate: $F(3,174)=35.06$, $p<0.0001$; Putamen: $F(3,174)=20.61$, $p<0.0001$; Globus Pallidus: $F(3,174)=13.53$, $p<0.0001$; Figure S9b). Specifically, we found that BG-cortical models exhibit highest prediction accuracy in contextual cortical representation (Caudate: Context vs. CxCO: $t(58)=6.37$, $p<0.0001$, Cohen's d=0.93; Context vs. CxSH: $t(58)=7.48$, $p<0.0001$, Cohen's d=1.10; Context vs. Decision: $t(58)=6.38$, $p<0.0001$, Cohen's d=0.63; Putamen: Context vs. CxCO: $t(58)=5.20$, $p=0.000016$, Cohen's d=0.78; Context vs. CxSH: $t(58)=5.18$, $p=0.000017$, Cohen's d=0.78; Context vs. Decision: $t(58)=4.16$, $p=0.0001$, Cohen's d=0.39; Caudate: Context vs. CxCO: $t(58)=4.09$, $p=0.0008$, Cohen's d=0.68; Context vs. CxSH: $t(58)=4.07$, $p=0.0009$, Cohen's d=0.66; Context vs. Decision: $t(58)=3.60$, $p=0.004$, Cohen's d=0.37), then followed by the second highest prediction accuracy in decision cortical representation (Caudate: CxCO vs. Decision: $t(58)=-2.78$, $p=0.04$, Cohen's d=-0.37; CxSH vs. Decision: $t(58)=-4.49$, $p=0.0002$, Cohen's d=-0.55; Putamen: CxCO vs. Decision: $t(58)=-3.43$, $p=0.007$, Cohen's d=-0.50; CxSH vs. Decision: $t(58)=-3.60$, $p=0.004$, Cohen's d=-0.49; Globus Pallidus: CxCO vs. Decision: $t(58)=-2.81$, $p=0.04$, Cohen's d=-0.44; CxSH vs. Decision: $t(58)=-2.84$, $p=0.04$, Cohen's d=-0.40). There was no significant difference between two relevant feature cortical representations.

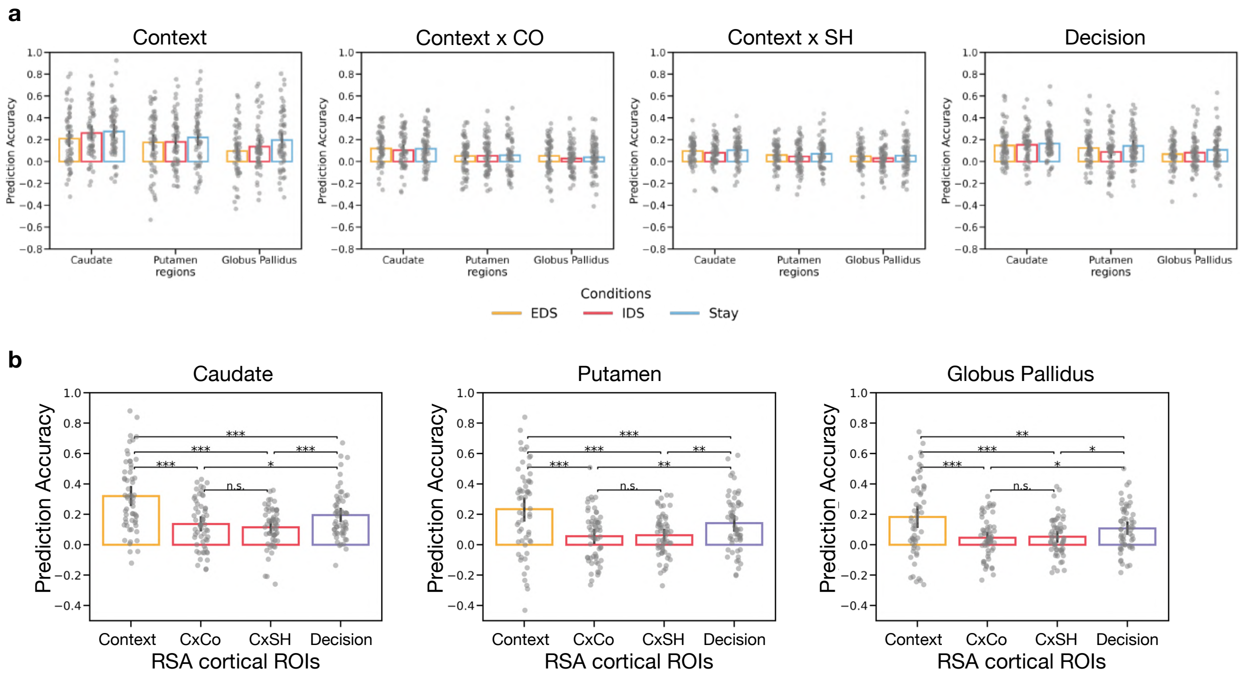


***S9. Voxel-to-voxel BG nuclei-cortical interaction model performances for predicting hierarchical task-switching conditions and cortical representations*** *(a) Model performance of three hierarchical task switching conditions when predicting different cortical representations; (b) Model performance of predicting different cortical representations. *p <0.05; **p <0.01; **p <0.001; n.s. not significant.*

**S10. Noise ceiling metric**

For the voxel-to-parcellation thalamocortical interaction model, we used a one-way repeated measure (rm) ANOVA to compare the split-half noise ceiling of voxel-to-parcellation thalamocortical interaction model for each task-switching condition (EDS, IDS and Stay; Table 12).

**Table 12: Noise ceiling results for voxel-to-parcellation thalamocortical interaction model**

| F-value (2, 116) | p-value | EDS (mean ± SD) | IDS (mean ± SD) | Stay (mean ± SD) |
|---|---|---|---|---|
| 0.10 | 0.37 | 0.67 ± 0.07 | 0.66 ± 0.07 | 0.67 ± 0.07 |

For the voxel-to-voxel thalamocortical interaction model, we used a one-way rmANOVA to compare the different conditions (EDS, IDS, Stay and All) within each cortical representation regions separately (Table 13).

**Table 13: Noise ceiling results for voxel-to-voxel thalamocortical interaction model**

| Region | F-value (3, 174) | p-value | EDS (mean ± SD) | IDS (mean ± SD) | Stay (mean ± SD) | All (mean ± SD) |
|---|---|---|---|---|---|---|
| Context | 2.26 | 0.08 | 0.60 ± 0.04 | 0.60 ± 0.04 | 0.60 ± 0.04 | 0.61 ± 0.04 |
| Context x Color | 4.41 | 0.01 | 0.59 ± 0.03 | 0.59 ± 0.03 | 0.60 ± 0.03 | 0.60 ± 0.03 |
| Context x Shape | 4.00 | 0.01 | 0.59 ± 0.02 | 0.59 ± 0.02 | 0.59 ± 0.02 | 0.59 ± 0.02 |
| Decision | 1.24 | 0.3 | 0.59 ± 0.03 | 0.59 ± 0.02 | 0.59 ± 0.02 | 0.60 ± 0.02 |

For the voxel-to-voxel caudate-cortical interaction model, we used a one-way rmANOVA to compare the different conditions (EDS, IDS, Stay and All) within each cortical representation regions separately (Table 14).

**Table 14: Noise ceiling results for voxel-to-voxel caudate-cortical interaction model**

| Region | F-value (3, 174) | p-value | EDS (mean ± SD) | IDS (mean ± SD) | Stay (mean ± SD) | All (mean ± SD) |
|---|---|---|---|---|---|---|
| Context | 2.74 | 0.04 | 0.59 ± 0.03 | 0.60 ± 0.03 | 0.59 ± 0.03 | 0.60 ± 0.03 |
| Context x Color | 1.95 | 0.12 | 0.58 ± 0.02 | 0.59 ± 0.02 | 0.59 ± 0.02 | 0.59 ± 0.02 |
| Context x Shape | 2.50 | 0.06 | 0.58 ± 0.01 | 0.58 ± 0.01 | 0.58 ± 0.01 | 0.58 ± 0.01 |
| Decision | 2.28 | 0.08 | 0.58 ± 0.01 | 0.57 ± 0.02 | 0.59 ± 0.01 | 0.59 ± 0.02 |

For the voxel-to-voxel putamen-cortical interaction model, we used a one-way rmANOVA to compare the different conditions (EDS, IDS, Stay and All) within each cortical representation regions separately (Table 15).

**Table 15: Noise ceiling results for voxel-to-voxel putamen-cortical interaction model**

| Region | F-value (3, 174) | p-value | EDS (mean ± SD) | IDS (mean ± SD) | Stay (mean ± SD) | All (mean ± SD) |
|---|---|---|---|---|---|---|
| Context | 3.64 | 0.01 | 0.59 ± 0.03 | 0.59 ± 0.03 | 0.60 ± 0.04 | 0.60 ± 0.03 |
| Context x Color | 3.40 | 0.02 | 0.58 ± 0.02 | 0.58 ± 0.02 | 0.59 ± 0.02 | 0.59 ± 0.02 |
| Context x Shape | 4.15 | 0.01 | 0.58 ± 0.01 | 0.58 ± 0.02 | 0.59 ± 0.02 | 0.59 ± 0.02 |
| Decision | 3.06 | 0.03 | 0.59 ± 0.02 | 0.58 ± 0.02 | 0.59 ± 0.02 | 0.59 ± 0.02 |

For the voxel-to-voxel globus pallidus-cortical interaction model, we used a one-way rmANOVA to compare the different conditions (EDS, IDS, Stay and All) within each cortical representation regions separately (Table 16).

**Table 16: Noise ceiling results for voxel-to-voxel globus pallidus-cortical interaction model**

| Region | F-value (3, 174) | p-value | EDS (mean ± SD) | IDS (mean ± SD) | Stay (mean ± SD) | All (mean ± SD) |
|---|---|---|---|---|---|---|
| Context | 2.92 | 0.04 | 0.59 ± 0.03 | 0.59 ± 0.03 | 0.60 ± 0.04 | 0.59 ± 0.03 |
| Context x Color | 0.39 | 0.76 | 0.58 ± 0.02 | 0.58 ± 0.02 | 0.58 ± 0.02 | 0.58 ± 0.02 |
| Context x Shape | 0.28 | 0.84 | 0.58 ± 0.01 | 0.58 ± 0.02 | 0.58 ± 0.023 | 0.58 ± 0.02 |
| Decision | 0.70 | 0.56 | 0.58 ± 0.02 | 0.58 ± 0.02 | 0.59 ± 0.02 | 0.59 ± 0.02 |
